# Plant resistance in different cell layers affects aphid probing and feeding behaviour during poor- and non-host interactions

**DOI:** 10.1101/372839

**Authors:** Carmen Escudero-Martinez, Daniel J. Leybourne, Jorunn I.B. Bos

**Affiliations:** Cell and Molecular Sciences, The James Hutton Institute, Invergowrie, Dundee, DD2 5DA, UK; Division of Plant Sciences, School of Life Sciences, University of Dundee, Dundee

**Keywords:** aphid, EPG analyses, nonhost, plant resistance, probing, stylet pathway

## Abstract

Aphids are phloem-feeding insects that cause economic losses to crops globally. Whilst aphid interactions with susceptible plants and partially resistant genotypes have been well characterised with regards to aphid probing and feeding behaviour, the interactions with non-natural host species are not well understood. Using aphid choice assays with the broad host range pest *Myzus persicae* and the cereal pest *Rhopalosiphum padi* we show that about 10% of aphids settle on non-/poor-host species over a 24h time period. We used the Electrical Penetration Graph technique to assess aphid probing and feeding behaviour during the non-/poor-host interactions. In the Arabidopsis non-host interaction with the cereal pest *R. padi* aphids were unable to reach and feed from the phloem, with resistance likely residing in the mesophyll cell layer. In the barley poor-host interaction with *M. persicae*, resistance is likely phloem-based as aphids were able to reach the phloem but ingestion was reduced compared with the host interaction. Overall our data suggests that plant resistance to aphids in non-host and poor-host interactions with these aphid species likely resides in different plant cell layers. Future work will take into account specific cell layers where resistances are based to dissect the underlying mechanisms and gain a better understanding of how we may improve crop resistance to aphids.

## Introduction

Aphids are important insect pests which cause significant yield losses to crops globally (Blackman R, 2000). There are approximately 5000 aphid species described and around 250 of these are important agricultural and horticultural pests which vary in their host range – the ability to successfully infest different plant species. This host range variation generally applies to secondary hosts during summer months, where aphid populations increase rapidly due to asexual reproduction (Moran, 1992). Whilst the majority of aphid species exhibit a limited host range, dedicated to few closely related plant species, some aphid species, like *Myzus persicae* Sulzer (green peach aphid), have an exceptionally broad host range which includes representatives from more than 40 plant families (Blackman R, 2000, Powell et al., 2006). The evolutionary drivers and molecular determinants of such exceptionally broad host ranges in aphids remain to be elucidated.

Host suitability relies on a number of factors, which could be based either at the plant surface or within plant tissues and cells (Powell et al., 2006). Prior to probing the leaf surface aphid behaviour can be influenced by a range of these factors including leaf colour, emitted volatile organic compounds and leaf surface components, such as epicuticular waxes or trichomes (Doring, 2014, Doring & Chittka, 2007, Neal et al., 1990). Regardless of whether the aphid encounters a host or non-host plant species their specialised mouthparts, known as stylets, are utilised to probe into the plant tissue (Escudero-Martinez et al., 2017, Jaouannet et al., 2015, Powell et al., 2006). This probing behaviour is associated with the transmission of important plant viruses during both host and non-host interactions (Debokx & Piron, 1990, Katis & Gibson, 1985, Powell et al., 2006, Verbeek et al., 2010) which can substantially reduce crop yields (Perry et al., 2000). During interactions with susceptible plant species the aphid stylets penetrate the plant epidermis and move through the plant tissue towards the vascular bundle. During this process the stylets probe into adjacent plant cells, and saliva is secreted both in the apoplast and into probed cells along the stylet-pathway (Tjallingii, 2006, Tjallingii & Esch, 1993). During compatible plant-aphid interactions the aphid stylets are able to successfully puncture the sieve-tube elements to facilitate ingestion of phloem sap (Tjallingii, 1995, Tjallingii, 2006).

The aphid stylet-pathway through the plant tissue has been well-characterised during interactions with susceptible plants using the Electrical Penetration Graph (EPG) technique. This technique uses an electrical circuit to connect the aphid to the plant via a series of electrical probes, allowing distinction between different phases of the stylet pathway from obtained electrical waveforms which correlate with the position of the aphid stylet within plant tissue in real-time (Prado & Tjallingii, 1994, Tjallingii, 1985a, Tjallingii, 1985b, Tjallingii & Esch, 1993). Briefly, the aphid is attached to an electrical probe with gold wire, and a copper electrode is placed into the soil to incorporate the plant into the electrical system. Both the plant and the aphid electrodes are attached to a data-logger, which is read by computational software and the whole set-up is contained in a grounded Faraday cage (Mclean & Kinsey, 1968, Tjallingii, 1978, Tjallingii, 1985a, Tjallingii, 1985b). Once the aphid probes the plant tissue the circuit closes and changes in electrical voltage are displayed as alternating waveforms which can be manually annotated using computational software and translated into time-series data (Tjallingii & Esch, 1993). The biological relevance of the different waveforms that are detected by the EPG technique have been extensively analysed (Prado & Tjallingii, 1994, Tjallingii, 1978, Tjallingii, 1985a, Tjallingii, 1985b). Waveforms associated with aphid probing are: waveform np, representing non-probing behaviour where the stylets are not in contact with the leaf surface; waveform C, which begins upon stylet penetration of leaf tissue and is correlated with the intercellular apoplastic stylet pathway located at the epidermis or the mesophyll cell layers; waveform pd, associated with piercing of a plant cell which leads to a signal potential drop; waveform F, which reflects stylet mechanical/penetration difficulties; and waveform E1e, which represents extracellular saliva secretion into plant tissues other than phloem. Waveforms associated with vascular interactions and which provide intricate information at the aphid feeding site are: waveform G, which represents aphids drinking from the xylem sap; waveform E1, which is linked to aphid salivation into phloem before ingestion; and waveform E2, which corresponds to phloem sap ingestion (Alvarez et al., 2006). A graphical representation of examples of these waveforms, alongside the stylet activity during each, is shown in Fig. 1.

**Figure 1.**
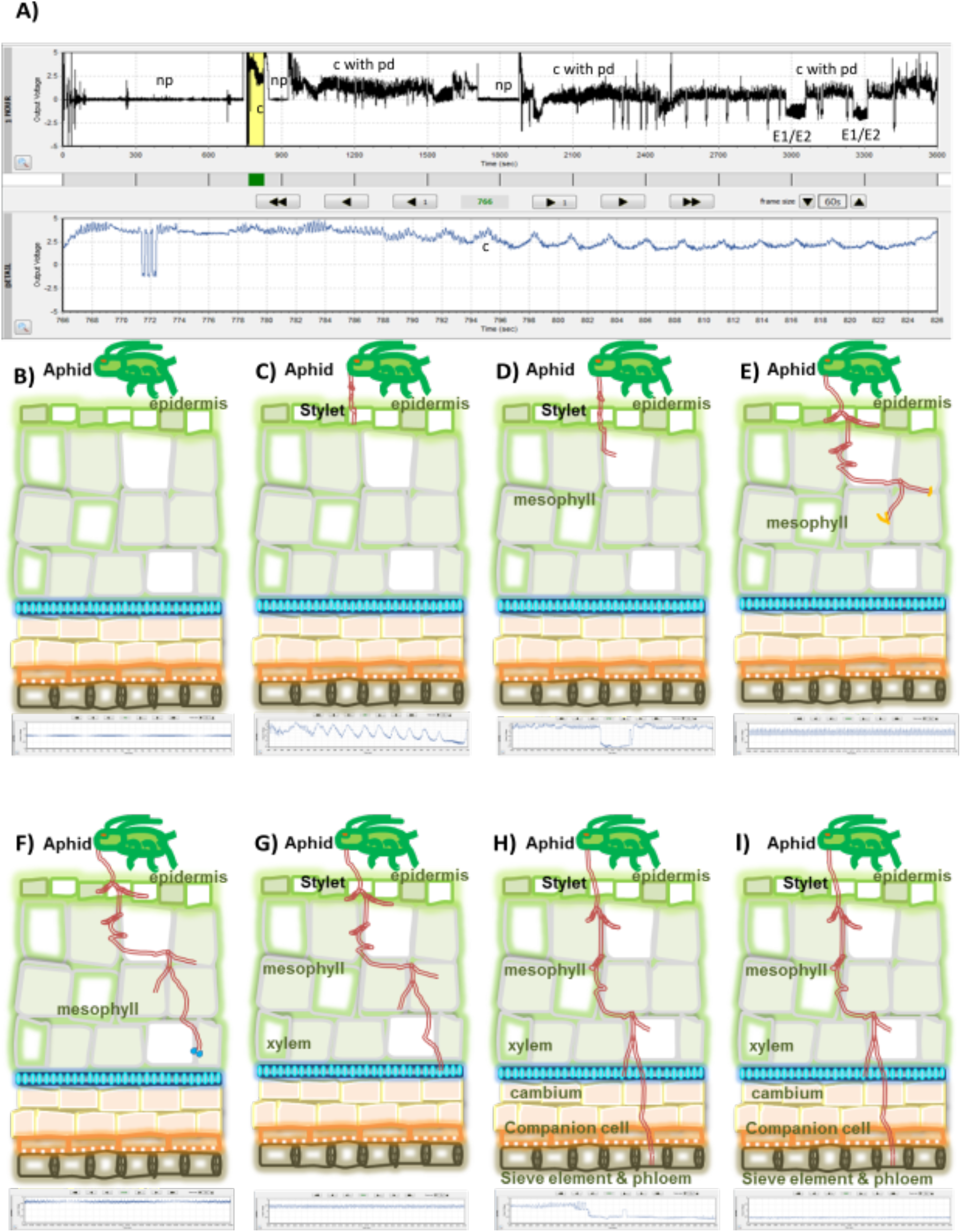
Graphical representation of aphid/stylet activities associated with each EPG waveform. (A) Example of aphid activity during np (non-probing) period, stylet is not in contact with leaf tissue. (B) Initiation of pathway (C) phase - aphid stylet pierces leaf epidermis, (C) Potential drop (pd) – aphid stylet penetrates adjacent plant cell (D) Stylet penetration difficulties (F phase) (E) Extracellular saliva secretion (E1e) phase – salivation into extracellular space. (F) Xylem ingestion (G phase) – stylet penetrates vascular xylem cells to initiate xylem drinking. (G) Salivation into phloem (E1 phase) – stylet penetrates sieve tube element and aphid initiates salivation into phloem sap. (H) Phloem ingestion (E2 phase) – aphid begins passive ingestion of phloem sap. Also includes sustained phloem ingestion (sE2 phase) - a period of phloem sap ingestion lasting > 10 mins.

Although the EPG technique has mainly been used to study aphid interactions with susceptible and (partially-)resistant genotypes of host plant species, it also represents a suitable tool to explore how aphids interact with plants which are not natural hosts, including non-host and poor-host species. Indeed, EPG analyses of *Brevicoryne brassicae* Linnaeus (cabbage aphid) on host Brassicaceae and non-host *Vicia faba* showed that this aphid species was unable to reach the phloem when feeding on the non-host *V. faba*, despite probing the leaf surface (Garbys & Pawluk, 1999). Also, epidermis and phloem factors contributed to resistance in different legume species to different pea aphid biotypes (Schwarzkopf et al., 2013). By characterising aphid probing and feeding behaviour across different aphid interactions with non-/poor-host species we aim to generate a better understanding of where associated resistance mechanisms reside. This in turn will facilitate important mechanistic studies to reveal the molecular determinants of plant immunity to aphids.

We previously showed that *M. persicae*, which is not a pest of barley, is able to feed and reproduce on this crop under controlled environment conditions, but to a lower extent than on a host species such as oil seed rape or Arabidopsis (Escudero-Martinez et al., 2017). On the contrary, *Rhopalosiphum padi* Linnaeus (bird cherry-oat aphid) is a pest of barley but is unable to feed from, and therefore survive, on Arabidopsis (Jaouannet et al., 2015). However, in both the *M. persicae*-barley poor-host interaction and the *R. padi*-Arabidopsis non-host interaction probing of the leaf surface takes place (Escudero-Martinez et al., 2017, Jaouannet et al., 2015). In line with our previous findings, choice assays showed that both aphid species will settle on and interact with non-/poor-host plant species if given a choice, with 10% of aphids found on non-/poor-hosts after 24h. Using EPG analyses of *M. persicae* and *R. padi* on Arabidopsis and barley we explored differences in aphid probing and feeding behaviour during non-/poor-host versus host interactions. We show that resistance in the non-/poor-host interactions can reside in different plant cell layers, suggesting complex mechanisms may underlie plant immunity to aphids.

## Materials and Methods

### Aphid rearing

*R. padi* (JHI-JB, genotype G) (J. et al., Thorpe et al., 2018) was maintained on *Hordeum vulgare* cv Optic and *M. persicae* (JHI_genotype O) was maintained on *Brassica napus* (oilseed rape). All aphid species used in the experiments were maintained in growth chambers under controlled conditions (18°C ± 2°C, 16 h of light).

### Plant growth

Barley plants (cv. Golden Promise) were pre-germinated in Petri dishes with wet filter paper for three days in the dark. Then, they were moved to a plant growth cabinet under controlled conditions and grown for 7 days (growth stage 1.10, determined using the staging key (Zadoks et al., 1974)) until the EPG experiments. *Arabidopsis thaliana* Col-0 plants were sown directly in soil; the seeds were stratified for 3 days at 4°C and placed in the growth cabinet for 4-5 weeks before use in experiments (growth stage 1.10 to 3.90, determined using the Boyes growth key (Boyes et al., 2001)). The cabinet conditions for Arabidopsis were 8 hours of light (125 μmol photons/m^2^.s), at 22 °C and 70% humidity. The cabinet conditions for barley were 8 hours of light (150 μmol photons/m2.s), at 20 °C (+-2°C).

### Aphid choice experiment

Aphid choice tests were devised to investigate the host plant preference of *R. padi* and *M. persicae*. Three choice test assays were developed: one using 50 *R. padi* aphids, a second using 50 *M. persicae* aphids, and a third using a mixed species population (25 *R. padi*, 25 *M. persicae*). For each assay, fifty aphids (mixed aged: 2^nd^ instar – apterous adult) were placed on a sheet of tissue paper and were placed in the centre of a Perspex cage halfway between two plants (one Arabidopsis, one barley). Aphids were 90 mm away from both plants and the two plants were 180 mm apart. Bamboo sticks served as bridges from the cage bottom (where the aphids were placed) to each plant, with additional bamboo sticks acting as bridges between the two plants, similar to the set-up used by Nowak and Komor (Nowak & Komor, 2010). Once the aphids were placed between the plants and the ladders were positioned, the cages were closed and the proportion of aphids present on the host, non-/poor-host, or which had not settled were scored three and 24 hours later. Choice assays were carried out in growth chambers under controlled conditions (18°C ± 2°C, 16 h of light).

Choice tests were carried out simultaneously in separate Perspex cages (440 mm x 340 mm x 390 mm). For each replicate the assignment of aphid mixture (*R. padi, M. persicae*, or mixed) to cage (1, 2, or 3) and the position (1 or 2) of Arabidposis and barley within each cage was randomly assigned. Seven replicates were collected for each aphid mixture. The proportion of aphids detected on each plant were modelled in response to plant type (Host, non-/poor-host, or not settled), aphid mixture (*R. padi*, *M. persicae*, mixed species), time-point (three hours and 24 hours) and all interactions using a linear mixed effects model. Cage and block were included as random factors, the model was simplified using manual backward stepwise model selection, and fitted-residual plots were observed at each stage to assess model suitability. Models were analysed using a χ^2^ Analysis of Deviance Test. Differences in the Least Squares Mean with Tukey correction for multiple comparison was used as a post-hoc test. Data were analysed in R Studio v. 1.0.143 running R v. 3.4.3 (R Core Team, 2017) with additional packages car v.2.1-4 (Weisberg & Fox, 2011), lme4 v.1.1-13, and lsmeans v.2.27-62 (Lenth, 2016).

### Electrical penetration graph (EPG) analyses

The probing and feeding behaviour of *R. padi* and *M. persicae* on different plant species was assessed using the Electrical Penetration Graph technique (Tjallingii, 1995) on a Giga-4 DC-EPG device with 1 Giga Ω resistance (EPG Systems, The Netherlands). We used a randomized block design for all EPG experiments performed here. Aphids were connected to a copper electrode with a golden wire (20 μm diameter), attached at the aphid dorsum and connected to the electrode with water-based silver glue. Aphids were lowered onto either an Arabidopsis or barley leaf approximately 1-1.5 hr after being removed from culture, depending on the treatment, and feeding behaviour was recorded over a 6h period. Three recordings were taken simultaneously. Each experiment was initiated between 10-12 am and the experiment was performed over a 6-month period, with 18 host and 17 non-host replicates for *R. padi* and 23 host and 28 poor-host replicates for *M. persicae*. Data were acquired using the Stylet+ D software package version v.01.28 and annotated manually using the Stylet+ A v.01.30 software (EPG-Systems, The Netherlands). Obtained waveforms were annotated with one of the following signals: no penetration (np), stylet penetration into the epidermal and mesophyll tissue (pathway/C phase), cellular punctures during the C phase (pd), watery salivation into sieve elements (E1), ingestion of phloem sap (E2), derailed stylet mechanics/stylet penetration difficulties (waveform F), xylem ingestion (waveform G), or extracellular saliva secretion into mesophyll (E1e) (Alvarez et al., 2006, Tjallingii, 1995). Annotated waveforms were converted into time-series data using the excel macro developed by Dr Schliephake (Julius Kühn-Institut); these converted parameters were used for statistical analysis. Parameters used for comparisons in these experiments are described by Giordanengo et al. (Giordanengo, 2014), and include total time of probing, number of probes, duration of phloem sap ingestion, and duration of xylem sap ingestion, a total of 97 parameters were measured. Statistical analyses were performed in R Studio running R v. 3.2.3. (R Core Team, 2017) using the Wilcoxon rank test, a significance threshold of 0.05 was used.

## Results

### Aphids preferentially settle on their host plant

We used aphid choice assays to examine the host plant preference of *Rhopalosiphum padi* and *Myzus persicae*. We monitored the settling behaviour of *R. padi* when provided with a choice between barley (host) and Arabidopsis (non-host), of *M. persicae* when provided with a choice between Arabidopsis (host) and barley (non-host), and of a mixed species population containing *R. padi* and *M. persicae*. The majority of aphids preferentially settled on the host plant, *c*. 50% of aphids settled on the host plant within three hours (Table 1; Fig. 2). The number of aphids that settled on the host plant increased to around 80% after 24 hours for all aphid populations assessed (*t* = −9.48; p = <0.001) with the number of unsettled aphids decreasing (*t* = 8.30; p = <0.001). However, approximately 10% of aphids were found on either the non-host or the poor-host plant at both time-points. No effect of aphid mixture was observed (Table 1), indicating that the presence of additional aphid species did not influence aphid behaviour.

**Table 1:** Statistical results of the choice test assay

| Response variable | Test Statistic (degrees of freedom) | p-value |
| --- | --- | --- |
| Plant Type | $X^2_{(2)} = 532.65$ | P = <0.001 |
| Aphid Mixture | $X^2_{(2)} = 0.01$ | P = 0.996 |
| Time-point | $X^2_{(1)} = 0.01$ | P = 0.949 |
| Plant Type x Aphid mixture | $X^2_{(4)} = 5.43$ | P = 0.245 |
| Plant Type x Time-Point | $X^2_{(2)} = 162.06$ | P = <0.001 |
| Aphid Mixture x Time-Point | $X^2_{(2)} = 0.01$ | P = 0.996 |
| Plant Type x Aphid Mixture x Time-Point | $X^2_{(4)} = 0.34$ | P = 0.986 |

**Figure 2.**
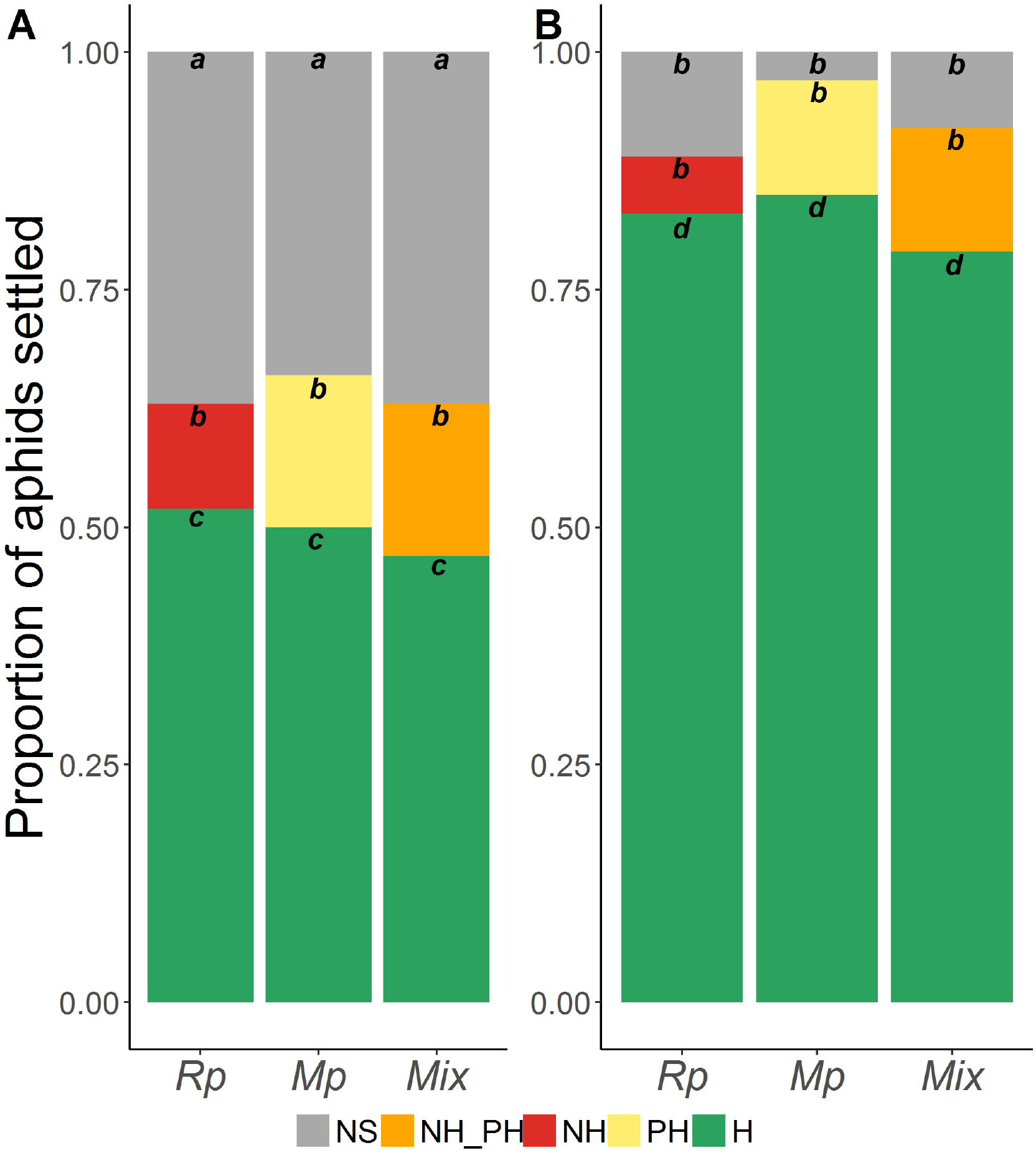
Stacked bar charts showing the settling behaviour of aphids in the choice experiment. (A) Aphid settling after three hours (B) Aphid settling after 24 hours Graphs show the mean proportion of aphids from the *R. padi* (Rp), *M. persicae* (Mp), and the mixed species population (Mix) which had settled on the host plant (H; green), the non-host plant (NH; red), the poor-host plant (PH; yellow), the non/poor-host plant (NH.PH; orange) or which has not settled (NS; grey). Letter under each bar indicate differences based on Least Squares Mean post-hoc analysis with Tukey correction.

### The Arabidopsis-*R. padi* non-host interaction is characterised by long no-probing periods and difficulties in locating the vascular tissues

We employed the Electrical Penetration Graph (EPG) technique to compare the feeding behaviour of *R. padi* on barley (host) with Arabidopsis (non-host) and of *M. persicae* on Arabidopsis (host) with barley (poor-host) over a six hour period in order to identify the tissue layers involved in non-host and poor-host resistance against aphids. We assessed 97 feeding parameters in total, 71 of these were altered during feeding on non/poor-host plants compared with feeding patterns on host plants (Table S1) with 26 parameters remaining unaffected (Table S2).

The majority of feeding parameters that differed between *R. padi* feeding on host compared with non-host plants were related to stylet probing of the plant tissue and interactions with the plant vasculature (Fig. 3). In general, probing parameters that differed for *R. padi* when interacting with non-host versus host plants were non-probing periods, number of stylet probes into plant tissue, and time spent in the epidermal/mesophyll cells (C phase) (Fig. 3A; Table S1).

**Figure 3.**
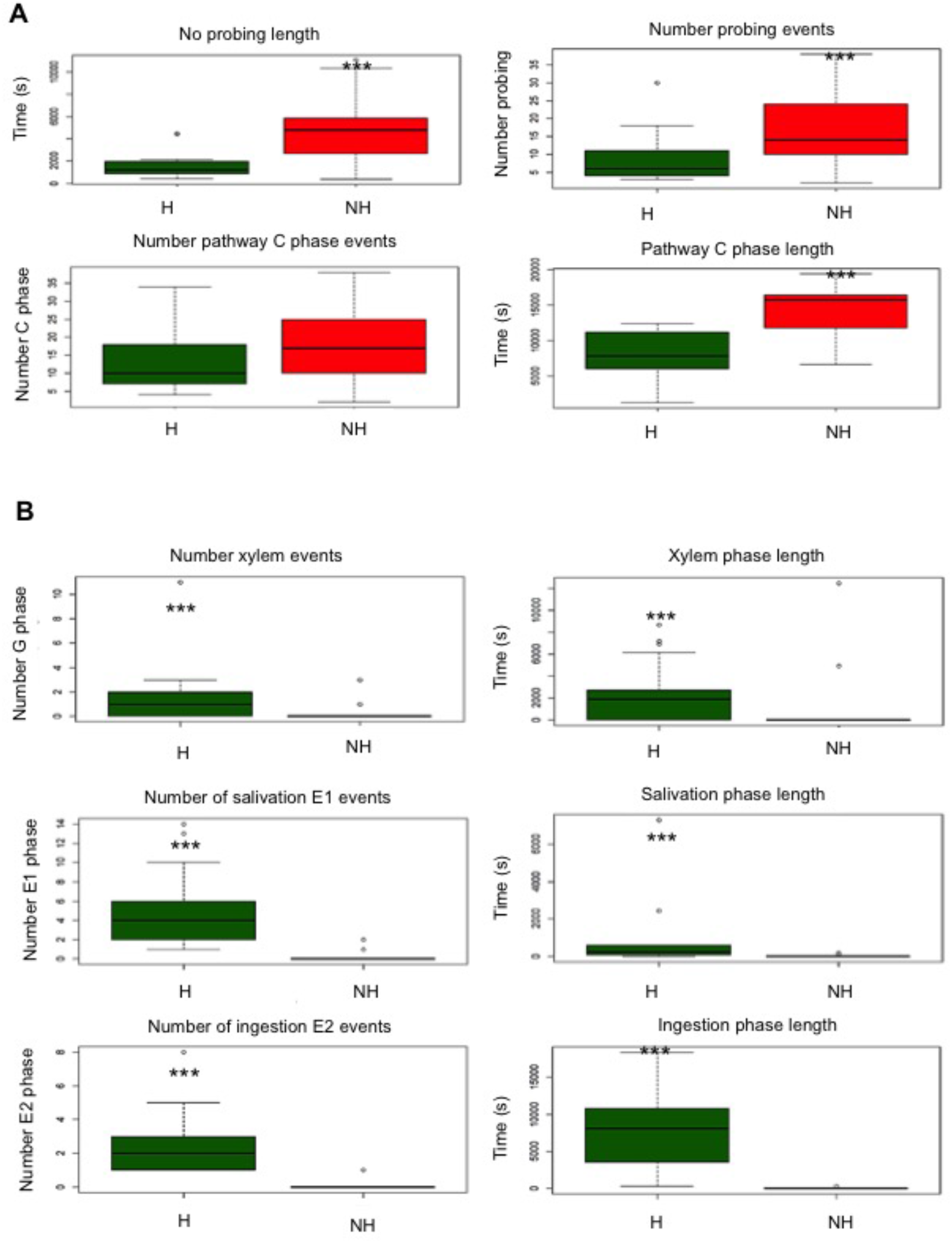
Box plots showing different EPG parameters associated with *Rhopalosiphum padi*-barley (host) and *Rhopalosiphum padi*-Arabidopsis (nonhost) interactions. (A) Probing-related parameters: total number of probing events, total length of no probing time, total number of pathway (C) phase events, total length of pathway (C) phase time. (B) Vascular-related parameters: number of xylem ingestion (G phase) events, total length of xylem ingestion, number of salivation (E1 phase) events where aphid saliva is secreted into phloem sap, total length of salivation (E1 phase), number of phloem sap ingestion (E2 phase) events and total length of phloem sap ingestion (E2 phase). Green boxes indicate the host (H) interaction and red boxes represent the non-host (NH) interaction. *R. padi* on host plants was replicated 18 times and *R. padi* on nonhost plants was replicated 17 times. Significant differences between interactions were assessed by Wilcoxon non-parametric t-test (*= p ≤0.05 and *** = p ≤0.01).

During non-host interactions with Arabidopsis, the total time the aphids were not probing plant tissue during the 6 h recording was 2.5 times greater (4889s) than the host interactions (1767s) (Fig. 3A; Table S1; W = 33.00; p = <0.001). However, the overall number of stylet probes into plant tissue was higher on non-host plants (18) than host plants (8) (Fig. 3A; Table S1; W = 52.50, p = 0.001). Although the total number of C phases (stylet activity at the epidermis/mesophyll, including a return to C phase following stylet interactions in the vasculature) was not significantly different between non-host and host interactions, the overall time spent in the epidermis/mesophyll (C phase) was over two times longer for the non-host (14128s) compared with host interactions (6237s) (Fig. 2A; Table S1; W = 37.00; p = <0.001).

All the vascular-related parameters (G, E1 salivation and E2 ingestion phases) measured for *R. padi* were significantly reduced during non-host interactions compared with host interactions (Fig. 3B; Table S1). This included a two-fold reduction in the number of xylem ingestion (G phase) events during the non-host interaction (0.24 times) compared with the host interaction (0.50 times) (Fig. 3B; Table S1; W = 2.28.50; p = 0.001) alongside a significant decrease in the total length of xylem ingestion, 1021s for non-host compared with 1483s for host plants (Fig. 3B; Table S1; W = 221.50; p = 0.003). We also observed significantly fewer salivation events (E1 phase) during the non-host interaction (0.18 events) compared with the host interaction (3.67 events; W = 282.00; p = <0.001), with salivation events five-fold shorter during the non-host interaction (18s) compared with the host interaction (93s) (Fig. 3B; Table S1; W = 278.00; p = <0.001). Ingestion of phloem sap (E2 phase) was rarely observed during the non-host interaction (0.06 times) compared with the host interaction (3 times; W = 285.00; p = <0.001), and the total duration of this ingestion period was greatly reduced on non-host plants (19s) compared with host plants (10030s, or 2.78 hours) (Fig. 3B; Table S1; W = 288.00; p = <0.001).

### The barley-*M. persicae* poor-host interaction is characterised by a lack of sustained phloem ingestion

The majority of feeding parameters that differed between *M. persicae* feeding on host compared with poor-host plants were primarily related to interactions within the plant vasculature, specifically a decrease in interactions with the phloem and an increase in interactions with the xylem (Fig. 4; Table S1). In general, this involved a decrease in the ability to locate the phloem and initiate ingestion of phloem sap. When feeding on poor-host plants there was a significant increase in the number of probes made into the plant tissue by aphids (19) compared with the number of probes made into host plants (16) (Fig. 3A; Table S1; W = 186.00; p = 0.024). However, the total length of time aphids probed into plant tissue, the number of pathway (C) phase events, and the total time spent within the pathway (C) phase was similar for the host and poor-host interactions (Fig. 4A)

Aphid stylet activities related to the vascular parameters (G – xylem, E1 – phloem salivation, and E2 – phloem ingestion) were different between host and poor-host interactions (Fig. 4B; Table S1). The number of times that *M. persicae* reached the xylem (G phase) during the poor-host interaction was higher (1.33 times; W = 133.50; p = <0.001) and the total time of xylem ingestion was longer (2321s; W = 142.50; p = <0.001) than during the host interaction, where aphids reached the xylem 0.30 times and spent a total of 691s ingesting xylem sap (Fig. 4B; Table S1). For the E1 salivation phase the number and duration of events was reduced during the poor-host interaction, 1.73 events (W = 5.28; p = <0.001) with a total length of time spent salivating into the phloem of 562s (W = 500.00; p = <0.001), compared with the host interaction (7 events with a time length of 652s) (Fig. 4B; Table S1).

**Figure 4:**
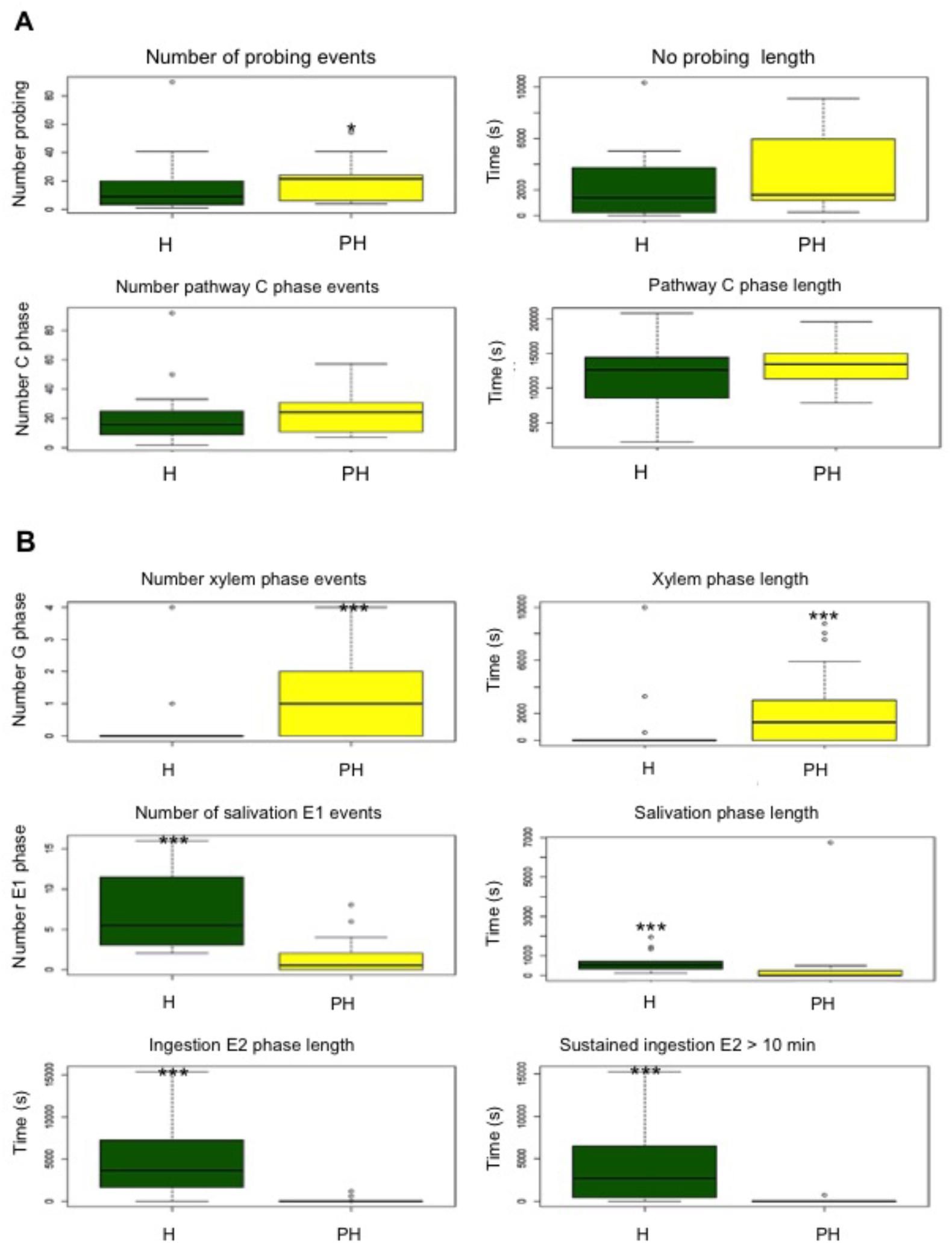
Box plots showing different EPG parameters in *Myzus persicae* interaction with a host (Arabidopsis) and a poor-host plant (barley). (A) Probing-related parameters: total number of probing events, total length of no probing time, total number of pathway (C) phase events, total length of pathway (C) phase time. (B) Vascular-related parameters: number of xylem ingestion (G phase) events, total length of xylem ingestion, number of salivation (E1 phase) events where aphid saliva is secreted into phloem sap, total length of salivation (E1 phase), total length of phloem sap ingestion (E2 phase) and total length of sustained phloem sap ingestion (sE2 phase). Green boxes indicate the host (H) interaction and yellow boxes represent the poorhost (PH)interaction. *M. persicae* on host plants was replicated 23 times and *M. persicae* on poor-host plants was replicated 28 times. Significant differences between interactions were assessed statistically by Wilcoxon non-parametric t-test (*= p ≤0.05 and *** = p ≤0.01).

*M. persicae* showed limited ingestion periods during the poor-host compared with host interactions. The number of E2 phases and their length was greatly reduced on poor-host plants, 0.53 events (W = 552.50; p = <0.001) with a 40-fold decrease in the total time spent ingesting phloem (126s; W = 573.50; p = <0.001), compared with host plants (5.7 events with a total length of 5064s) (Fig. 4B; Table S1). Moreover, on the poor-host sustained phloem ingestion was severely lacking, and aphids spent only 49s in the E2 ingestion phase on poor-host plants (W = 520.00; p= <0.001) with events being nearly absent, 0.07 events (W = 515.00; p = <0.001). In contrast, aphids spent 4322s in the E2 sustained ingestion phase on host plants over 2.1 events during the 6h recording (Fig. 4B; Table 1). Therefore, the *M. persicae* poor-host interaction features substantially reduced phloem ingestion.

## Discussion

The overall aim of this study was to gain insight into where resistances against aphids may reside within the plant tissue during host versus non/poor-host interactions by analysing aphid probing and feeding behaviour. We showed that when given a choice aphids do interact with non-/poor-host plants under controlled conditions, and we further explored these interactions using EPG analyses. Common features of the non-host and poor-host interactions were an increased number of probes and longer no-probing periods. Importantly, our data showed differences between *R. padi* and *M. persicae* probing and feeding behaviour on the non-/poor-host plants. During the *R. padi*-Arabidopsis (non-host) interaction the aphids only occasionally reached the vascular tissues. On the contrary, during the *M. persicae*-barley interaction (poor-host) aphids successfully reached the vascular tissue and could ingest xylem and phloem, however prolonged periods of phloem ingestion were inhibited. Based on the data generated here for *M. persicae* and *R. padi* we propose a model wherein poor- and non-host plant resistances against these aphid species may reside within the phloem and mesophyll cell layers, respectively (Fig. 5).

**Figure 5.**
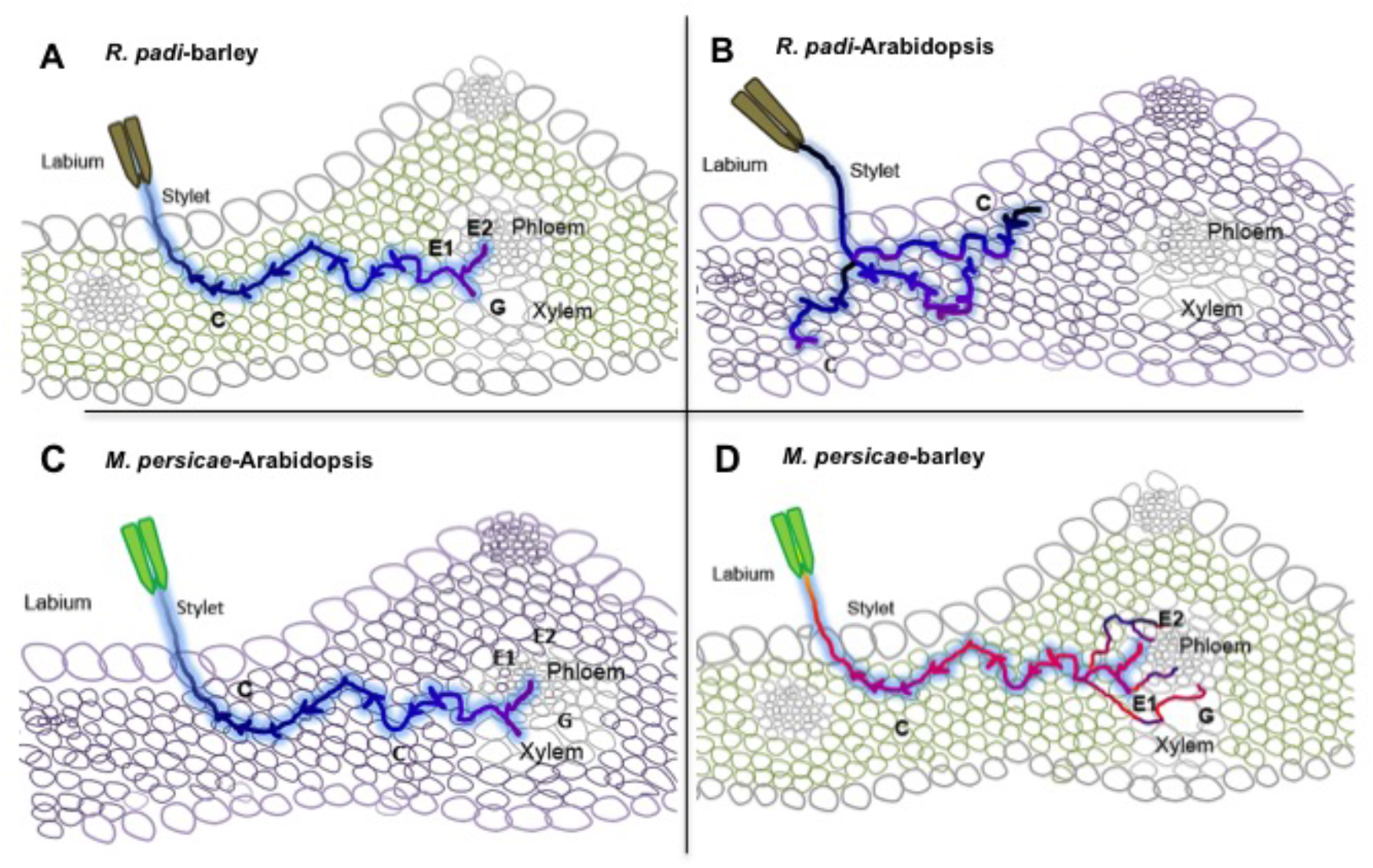
Model showing *R. padi* and *M. persicae* probing and feeding during host, poor-host and non-host plant interactions. (A) During the host interaction (*R. padi*-barley), the aphids will probe the epidermal and mesophyll cells (pathway C phase), then will drink from the xylem or salivate and feed from the phloem, with feeding lasting for hours. (B) During the non-host interaction (*R. padi*-Arabidopsis), the aphids will spend a long time not probing, and when probing eventually occurs the aphids remain in stylet pathway phase (in epidermis and mesophyll cell layers) most of the time and only occasionally will reach the vascular tissue, either xylem or phloem. No sustained ingestion of phloem sap takes place. (C) During the host interaction (*M. persicae*-Arabidopsis), the aphids will probe the epidermal and mesophyll cells (pathway C phase), then will drink from the xylem or salivate and feed from the phloem, with feeding taking place for hours. (D) During the poor-host interaction (*M. persicae*-barley), the aphids show increased probing compared to the host interaction, while the stylet pathway phase (in epidermis and mesophyll cell layers) is similar to the interaction with the host plant. At the vascular level, long periods of time will be spent in the xylem, and eventually aphid will reach the phloem, salivate and ingest phloem sap. However, contrary to the host interaction, no sustained (>10 minutes) ingestion of phloem sap takes place.

During the *R. padi*-barley interaction (host interaction) the aphids spend less time probing and in the pathway (C) phase and readily reach the phloem where salivation and phloem sap ingestion occurs for several hours (Fig. 5A). Occasionally, aphids ingest xylem, which is thought to be important in coping with osmotic effects associated with ingestion of large amounts of phloem sap (Pompon et al., 2010, Spiller et al., 1990). In contrast, during the *R. padi* – Arabidopsis interaction (non-host interaction) aphids exhibit altered probing behaviour, including an increase in the number of plant probes alongside a decrease in the total time probing into plant tissue. Additionally, *R. padi* shows an extended stylet pathway phase, and only rarely does the aphid reach the Arabidopsis phloem or xylem (Fig. 5B). On the occasions where the *R. padi* stylets reach the vascular tissue during non-host interactions the ingestion of phloem and xylem sap is ineffective, in line with this aphid being unable to survive on Arabidopsis (Jaouannet et al., 2015).

Interestingly, *R. padi* spent less time probing into plant tissue during the non-host interaction. However, during these probes aphids spent an increased time interacting with the mesophyll tissue during the non-host interaction than the host interaction, including an increase in the total time spent in the pathway (C) phase. This indicates that non-host resistance could potentially reside in the mesophyll tissue as the aphids struggled to probe beyond this layer and access to the vascular tissue was limited (Fig. 5B), as further indicated by the increased time required for aphids to reach the phloem during non/poor-host interactions compared with the host interactions. Further research will be needed to further understand the mechanisms underlying Arabidopsis non-host resistance to *R. padi*, and to investigate the potential involvement of specific recognition receptors within the mesophyll cell layer. Interestingly, the NADPH oxidase *AtRbohF*, involved in ROS (Reactive Oxygen Species) production, a member of the *LEA* (*Late Embryogenesis Abundant*) family, implicated in abiotic and biotic stress, as well as the *VSP1* (*Vegetative Storage Protein 1*), which is activated by jasmonate signalling, contribute to Arabidopsis non-host resistance against *R. padi* (Jaouannet et al., 2015). Whether these genes act within the mesophyll cell layer to activate defences against aphids remains to be determined.

The *M. persicae*-Arabidopsis (host) interaction, features short probing and pathway times, and prolonged salivation and ingestion once the phloem is reached, as well as occasional xylem drinking (Fig. 5C). In contrast, during the *M. persicae*-barley interaction (poor-host interaction) aphids show increased probing but spend a similar time in the stylet pathway phase as aphids on host Arabidopsis plants. The main differences between the Arabidopsis (host) and barley (poor-host) interactions with *M. persicae* are reduced salivation in the phloem and relatively short periods of phloem ingestion (less than 10 minutes) on barley (Fig. 5C and 5D). It is likely that this reduced phloem sap ingestion is responsible for the reduced *M. persicae* performance on barley (Escudero-Martinez et al., 2017, Ramirez & Niemeyer, 2000). It is possible that *M. persicae* attempts to compensate for this reduced ingestion of phloem sap with increased xylem drinking, in line with the observation that aphid starvation increases the xylem phase (Fig. 5D) (Ramirez & Niemeyer, 2000).

Phloem resistance factors are related to the E1 salivation and E2 ingestion parameters, and in particular ingestion phases shorter than 10 minutes (Alvarez et al., 2006, Prado & Tjallingii, 1997). Phloem-mediated defences against aphids include the occlusion of sieve elements, which prevents aphids from ingesting phloem sap (Dreyer & Campbell, 1987, Medina-Ortega & Walker, 2015, Will & van Bel, 2006). This phloem occlusion occurs upon callose deposition and formation of P-protein plugs. The latter is thought to seal off the phloem upon damage and/or to block the aphid food canal (Tjallingii, 2006, Will & van Bel, 2006). Interestingly, PAD4 was found to be a component of phloem-based immunity against *M. persicae* in Arabidopsis (Pegadaraju et al., 2007). However, no barley PAD4 (MLOC_1340) or PAD4-related genes were up-regulated during the barley-*M. persicae* interaction (Escudero-Martinez et al., 2017). However, our previous transcriptome analyses showed induction of a barley gene encoding Phloem Protein 2-like (PP2), which is a phloem specific lectin, with the induction being most pronounced during the barley-*M. persicae* interaction (Escudero-Martinez et al., 2017). Lectins have carbohydrate-binding properties and function in cell communication, development, and plant defence (Bellande et al., 2017). PP2 is a lectin highly abundant in the phloem and accumulates in damaged phloem sieve pores to form protective plugs (Read & Northcote, 1983). Overexpression of *AtPP2* in Arabidopsis leads to reduced *M. persicae* feeding suggesting PP2 may contribute to defences against aphids (Zhang et al., 2011), possibly by interfering with aphid digestion in the midgut (Kehr, 2006). The very infrequent phloem sap ingestion we observed might reflect a rejection of the sieve element, possibly due to the presence of a deterrent factor in the phloem sap (Mayoral et al., 1996). Indeed, lectins, including PP2-like proteins, have been shown to have deterrent activities and insecticidal activities against *M. persicae* (Jaber et al., 2010, Sauvion et al., 1996, Zhang et al., 2011). Whether barley phloem-lectins like PP2 indeed contribute to phloem-based defences of barley against *M. persicae* needs to be further tested.

It is important to note that the EPG experimental set-up was of a no-choice nature (i.e. aphids were placed on the plants) and that additional plant resistance components that affect aphid choice may play a role in the interactions studied here (Escudero-Martinez et al., 2017, Powell et al., 2006). For example, we previously showed that the black cherry aphid (*Myzus cerasi* Fabricius), which infests cherry trees as well as several herbaceous plants, displays only limited probing on non-host barley plants, and does not settle on barley leaves (Escudero-Martinez et al., 2017), pointing to a potential role of barley defences that act at the pre-probing level against this aphid species (Nottingham et al., 1991). In addition, some plant induced volatile compounds have been reported to be repellent to aphid pests and attractants of their natural enemies (Dreyer & Jones, 1981, Mallinger et al., 2011, Turlings & Ton, 2006).

With limited genetic crop resistance available against aphids, identifying the determinants of non/poor-host resistance is an important area of research that may help the development novel crop protection strategies. Using a detailed assessment of aphid probing and feeding behaviour on different natural host and non-host species we show that resistances may reside in different cell layers depending on the plant species-aphid species interaction.

## Acknowledgements

Dr. Freddy Tjallingii (EPG Systems, The Netherlands), Professor Alberto Fereres (CSIC, Spain) and Professor Gregory Walker (University of California, Riverside, USA) for providing EPG training, and additional thanks to Dr Tjallingii for helpful comments on non-host EPG waveforms. We also thank Dr. Nicholas Birch (The James Hutton Institute) for allowing us to use the EPG equipment. This work was support by the European Research Council (310190-APHIDHOST to JIBB), and the James Hutton Institute and Universities of Aberdeen and Dundee through a Scottish Food Security Alliance (Crops) PhD studentship to DJL.

## Author contributions

JIBB, CEM and DJL conceived and designed the experiments, CEM and DJL performed the experiments, JIBB, CEM and DJL analysed the data, JIBB and CEM wrote the manuscript with input from DJL. All authors read and approved the final manuscript.

## Supplementary Data

**Table S1**. Results for all obtained Electrical penetration graph (EPG) parameters which were significantly different between host and non/poor-host feeding. Table displays the EPG parameter assessed, a description of the parameter, and the plant tissue layer involved. Results displayed are the mean and standard deviation (SD) for each aphid-plant combination for each parameter alongside the Wilcoxon test statistic (W value) and p value for each pairwise host vs non/poor host comparison. p values in bold represent values significantly different in both host vs non-host and host vs poor-host interactions, italicised p values represent parameters which only differed in one combination. Average and standard deviation of the 97 electrical EPG parameters calculated for *R. padi* host (Rp_Hv) and non-host (Rp_At). Average and standard deviation of the 97 electrical EPG parameters calculated for *M. persicae* host (Mp_At) and poor-host (Mp_Hv). Calculations were made with summary statistics in Rstudio. The EPG list of parameters was taken from EPG systems: www.epgsystems.eu/files/List%20EPG%20variables.xls

**Table S2:** Results for all obtained Electrical penetration graph (EPG) parameters which were not significantly different between host and non/poor-host feeding. Table displays the EPG parameter assessed, a description of the parameter, and the plant tissue layer involved. Results displayed are the mean and standard deviation (SD) for each aphid-plant combination for each parameter alongside the Wilcoxon test statistic (W value) and p value for each pairwise host vs non/poor host comparison. p values in bold represent values significantly different in both host vs non-host and host vs poor-host interactions, italicised p values represent parameters which only differed in one combination. Average and standard deviation of the 26 electrical EPG parameters calculated for *R. padi* host (Rp_Hv) and non-host (Rp_At). Average and standard deviation of the 97 electrical EPG parameters calculated for *M. persicae* host (Mp_At) and poor-host (Mp_Hv). Calculations were made with summary statistics in Rstudio. The EPG list of parameters was taken from EPG systems: www.epgsystems.eu/files/List%20EPG%20variables.xls

